# Recommendations to address uncertainties in environmental risk assessment using toxicokinetics-toxicodynamics models

**DOI:** 10.1101/356469

**Authors:** Virgile Baudrot, Sandrine Charles

## Abstract

Providing reliable environmental quality standards (EQSs) is a challenging issue in environmental risk assessment (ERA). These EQSs are derived from toxicity endpoints estimated from dose-response models to identify and characterize the environmental hazard of chemical compounds such as those released by human activities. These toxicity endpoints include the classical *x*% effect/lethal concentrations at a specific time *t* (*EC/LC*(*x*, *t*)) and the new multiplication factors applied to environmental exposure profiles leading to *x*% effect reduction at a specific time *t* (*MF*(*x*, *t*), or denoted *LP*(*x*, *t*) by the EFSA). However, classical dose-response models used to estimate toxicity endpoints have some weaknesses, such as their dependency on observation time points, which are likely to differ between species (e.g., experiment duration). Furthermore, real-world exposure profiles are rarely constant over time, which makes the use of classical dose-response models difficult and compromises the derivation of *MF*(*x*, *t*). When dealing with survival or immobility toxicity test data, these issues can be overcome with the use of the general unified threshold model of survival (GUTS), a toxicokinetics-toxicodynamics (TKTD) model that provides an explicit framework to analyse both time- and concentration-dependent data sets as well as obtain a mechanistic derivation of *EC/LC*(*x*, *t*) and *MF*(*x*, *t*) regardless of x and at any time t of interest. In addition, the assessment of a risk is inherently built upon probability distributions, such that the next critical step for ERA is to characterize the uncertainties of toxicity endpoints and, consequently, those of EQSs. With this perspective, we investigated the use of a Bayesian framework to obtain the uncertainties from the calibration process and to propagate them to model predictions, including *LC*(*x*, *t*) and *MF*(*x*, *t*) derivations. We also explored the mathematical properties of *LC*(*x*, *t*) and *MF*(*x*, *t*) as well as the impact of different experimental designs to provide some recommendations for a robust derivation of toxicity endpoints leading to reliable EQSs: avoid computing *LC*(*x*, *t*) and *MF*(*x*, *t*) for extreme *x* values (0 or 100%), where uncertainty is maximal; compute *MF*(*x*, *t*) after a long period of time to take depuration time into account and test survival under few correlated and uncorrelated pulses of the contaminant in terms of depuration.

## Introduction

Assessing the environmental risk of chemical compounds requires the definition of environmental quality standards (EQSs). EQS are based on several calculations depending on the context and institutions such as predicted-no-effect concentrations (PNECs) [19] and specific concentration limits (SCLs) [17]. Specifically, the derivation of EQSs results from a combination of assessment factors with toxicity endpoints mainly estimated from measured exposure responses of a set of target species to a certain chemical compound [17, 19, 27, 37]. Estimating reliable toxicity endpoints is challenging and very controversial [28, 32]. Currently, the first step of environmental risk assessment (ERA) is the hazard identification of acute effects, which consists of fitting classical dose-response models to quantitative toxicity test data. For acute effect assessment, such data are collected from standard toxicity tests, from which the 50% lethal or effective concentration (*LC*_50_ or *EC*_50_, respectively) is generally estimated at the end of the exposure period, meaning that not all observations are used. In addition, classical dose-response models implicitly assume that the exposure concentration remains constant throughout the experiment, which makes it difficult to extrapolate the results to more realistic scenarios with time-variable exposure profiles combining different heights, widths and frequencies of contaminant pulses [7, 13, 28, 35].

To overcome this limitation at the organism level, the use of mechanistic models, such as toxicokinetics-toxicodynamics (TKTD) models, is now promoted to describe the effects of a substance of interest by integrating the dynamics of the exposure [19, 26, 29]. Indeed, TKTD models appear highly advantageous in terms of gaining a mechanistic understanding of the chemical mode of action, deriving time-independent parameters, interpreting time-varying exposure and making predictions under untested conditions [7, 29]. Another advantage of TKTD models for ERA is the possible calculation of lethal concentrations for any *x*% of the population at any given exposure duration *t*, denoted *LC*(*x*, *t*). Furthermore, from time-variable concentration profiles observed in the environment, it is possible to estimate a margin of safety such as the exposure multiplication factor *MF* (*x*, *t*), leading to any *x*% effect reduction due to the contaminant at any time *t* [7, 20] (also called the lethal profile and denoted *LP* (*x*, *t*) by [20]).

When focusing on the survival rate of individuals, the general unified threshold model of survival (GUTS) has been proposed to unify the majority of TKTD survival models [29]. In the present paper, we consider the two most used derivations, namely, the stochastic death (GUTS-RED-SD) and individual tolerance (GUTS-RED-IT) models. The GUTS-RED-SD model assumes that all individuals are identically sensitive to the chemical substance by sharing a common internal threshold concentration and that mortality is a stochastic process once this threshold is reached. In contrast, the GUTS-RED-IT model is based on the critical body residue (CBR) approach, which assumes that individuals differ in their thresholds, following a probability distribution, and die as soon as the internal concentration reaches the individual-specific threshold [29]. The robustness of GUTS models in calibration and prediction has been widely demonstrated, with little difference between GUTS-RED-SD and GUTS-RED-IT models [7, 10, 30]. Sensitivity analysis of toxicity endpoints derived from GUTS models, such as *LC*(*x*, *t*) and *MF* (*x*, *t*), has also been investigated [7, 10], but the question of how uncertainties are propagated is still under-studied.

Quantifying uncertainties or levels of confidence associated with toxicity endpoints is undoubtedly a way to improve trust in risk predictors and to avoid decisions that could increase rather than decrease the risk [12, 14, 24]. The Bayesian framework has many advantages for dealing with uncertainties since the distribution of parameters and thus their uncertainties is embedded in the inference process [36]. While the construction of priors on model parameters can be seen as subjective [21], it provides added value by taking advantage of information from the experimental design [10, 15]. Consequently, coupling TKTD models with Bayesian inference allows one to estimate the probability distribution of toxicity endpoints and any other predictions coming from the mechanistic (TKTD) model by taking into account all the constraints resulting from the experimental design. Moreover, Bayesian inference, which is particularly efficient with GUTS models [10, 15], can also be used to optimize the experimental design by quantifying the gain in knowledge from priors to posteriors [1]. Finally, Bayesian inference is tailored for decision making as it provides assessors with a range of values rather than a single point, which is particularly valuable in risk assessment [21, 24].

In the present study, we explore how scrutinizing uncertainties helps provide recommendations for experimental design and the characteristics of toxicity endpoints used in EQSs while maximizing their reliability. We first give an overview of TKTD models, with a focus on the GUTS [29] to derive EQS explicite equations. We then illustrate how to handle GUTS models within the R package *morse* [8] with five example data sets. Then, we explore how a variety of experimental designs influence the uncertainties in derived *LC*(*x*, *t*) and *MF* (*x*, *t*). Finally, we provide a set of recommendations on the use of TKTD models for ERA based on their added value and the way the uncertainty may be handled under a Bayesian framework.

## Material and methods

### Data from experimental toxicity tests

We used experimental toxicity data sets described in [5] and [33] testing the effect of five chemical compounds (carbendazim, cypermethrin, dimethoate, malathion and propiconazole) on the survival rate of the amphipod crustacean *Gammarus pulex*. Two experiments were performed for each compound, one exposing *G. pulex* to constant concentrations and the other exposing *G. pulex* to time-variable concentrations (see Table 1). In the constant exposure experiments, *G. pulex* was exposed to eight concentrations for four days. In the time-variable exposure experiments, *G. pulex* was exposed to two different pulse profiles consisting of two one-day exposure pulses with either a short or long interval between them.

### GUTS modelling

In this section, we detail the mathematical equations of GUTS models describing the survival rate over time of organisms exposed to a profile of concentrations of a single chemical product. All other possible derivations of GUTS models are fully described in [29, 30]. Here, we provide a summary of GUTS-RED-SD and GUTS-RED-IT reduced models to introduce notations and equations relevant for mathematical derivation of explicit formulations of the *x*% lethal concentration at time *t*, denoted *LC*(*x*, *t*), and of the multiplication factor leading to *x*% mortality at time *t*, denoted *MF* (*x*, *t*).

**Table 1.** Characteristics of data sets used in the manuscript. “Profile type” is the type of exposure profile (constant or time-variable), “Data points” refers to the number of data points in the data set, “Nbr profiles” is the number of profiles in the data set, “*N*_*init*_” is the initial number of individuals in the profile, “Nbr days” is the number of days for each experiment, and “Time points per profile” is the number of observation time points for each time series (each constant profile consisted of 5 time points).

| Product | Profile type | Data points | Nbr profiles | $N_{init}$ | Nbr days | Time points per profile |
| --- | --- | --- | --- | --- | --- | --- |
| carbendazim | constant | 40 | 8 | 20 | 4 | 5 |
| cypermethrin | constant | 40 | 8 | 20 | 4 | 5 |
| dimethoate | constant | 40 | 8 | 20 | 4 | 5 |
| malathion | constant | 40 | 8 | 20 | 4 | 5 |
| propiconazole | constant | 40 | 8 | 20 | 4 | 5 |
| carbendazim | variable | 51 | 4 | 80 | 10 | [8, 14, 16, 13] |
| cypermethrin | variable | 61 | 4 | 80 | 10 | [10, 18, 18, 15] |
| dimethoate | variable | 58 | 4 | 80 | 10 | [10, 16, 17, 15] |
| malathion | variable | 70 | 2 | 70 | 22 | [35, 35] |
| propiconazole | variable | 74 | 4 | 70 | 10 | [11, 21, 21, 21] |

### Toxicokinetics

We define *C*_*w*_(*t*) as the external concentration of a chemical product, which can be variable over time. As there is no measure of internal concentration, we use the scaled internal concentration, denoted *D*_*w*_(*t*), which is therefore a latent variable described by the toxicokinetics part of the model as follows:

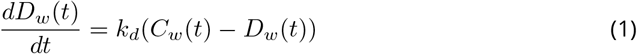

where *k*_*d*_ [*time*^−1^] is the dominant rate constant, corresponding to the slowest compensating process dominating the overall dynamics of toxicity.

As we assume that the internal concentration equals 0 at *t* = 0, the explicit formulation for constant concentration profiles is given by

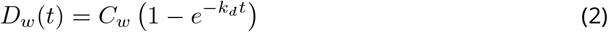

An explicit expression for time-variable exposure profiles is provided in the Supplementary Material as it can be useful for implementation but not for the mathematical calculus presented below. The GUTS-RED-SD and GUTS-RED-IT models are based on the same model for the scaled internal concentration. These models do not differ in the TK part but do differ in the TD part describing the death mechanism.

From the toxicokinetics equation (2), we can easily compute the *x*% depuration time *DRT*_*x*_, that is, the period of time after a pulse leading to an *x*% reduction in the scaled internal concentration:

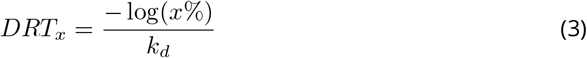

While GUTS-RED-SD and GUTS-RED-IT models have the same toxicokinetic equation (1), the *DRT*_*x*_ likely differs between them since the meaning of damage depends on the toxicodynamic equations, which are different.

### Toxicodynamics

The GUTS-RED-SD model supposes that all the organisms have the same internal threshold concentration, denoted *z* [*mol.L*^−1^], and that once this concentration threshold is exceeded, the instantaneous probability of death, denoted *h*(*t*), increases linearly with the internal concentration. The mathematical equation is

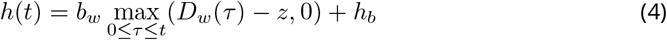

where *b*_*w*_ [*L.mol.time*^−1^] is the killing rate and *h*_*b*_ [*time*^−1^] is the background mortality rate.

Then, the survival probability over time under the GUTS-RED-SD model is given by

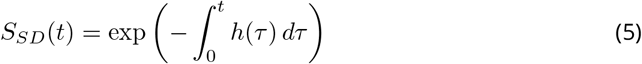

The GUTS-RED-IT model supposes that the threshold concentration is distributed among organisms and that death is immediate as soon as this threshold is reached. The probability of death at the maximal internal concentration with background mortality *h*_*b*_ is given by

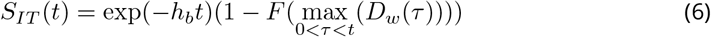

Assuming a log-logistic function, we get 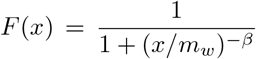, with the median *m*_*w*_ [*mol.L*^−1^] and shape *β* of the threshold distribution, which gives

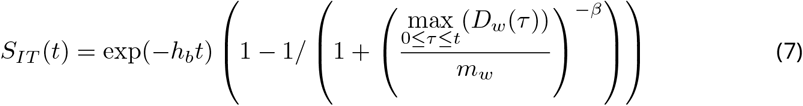

### Implementation and Bayesian inference

GUTS models were implemented within a Bayesian framework with *JAGS* [34] by using the R package *morse* [8]. The Bayesian inference methods, choice of priors and parameterisation of the MCMC process have previously been fully explained [8, 10, 15]. The joint posterior distribution of parameters was used to predict survival curves under tested and untested exposure profiles, to calculate *LC*(*x*, *t*) and *MF* (*x*, *t*), and to compute goodness-of-fit measures (see hereinafter). The use of the joint posterior distribution allowed us to quantify the uncertainty around all these predictions; therefore, their medians and 95% credible intervals were computed as follows: under a specific exposure profile, we simulated the survival rate over time for every joint posterior parameter set; then, at each time point of the time series, we computed 0.5, 0.025 and 0.975 quantiles, thus providing medians and 95% limits.

### Measures of model robustness

Modelling is always associated with testing robustness: not only the robustness in fitting data used for calibration but also the robustness in generating predictions with new data [25]. To evaluate the robustness of estimations and predictions with the two GUTS models, we calculated their statistical properties by means of the normalized root mean square error (NRMSE), the posterior predictive check (PPC), the Watanabe-Akaike information criterion and leave-one-out cross-validation (LOO-CV) [23].

### Normalized root mean square error

The root mean square error (RMSE) allows one to characterize the difference between observations and predictions from the posterior distribution. With *N* observations and *y_i,obs_* observed individuals (*i* ∈ {1, …, *N*}), for each estimation *y*.,_*j*_ of the Markov chain of size *M* (*j* ∈ {1, …, *M*}) resulting from the Bayesian inference, we can define the *RMSE*_*j*_ as

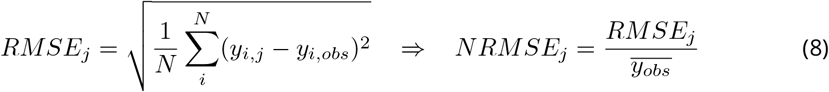

where the normalized RMSE (NRMSE) is given by dividing RMSE by the mean of the observations, denoted 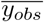. We then have the distribution of the NRMSE, from which we can obtain the median and the 95% credible interval, as presented in Table 2.

### Posterior predictive check (PPC)

The posterior predictive check consists of comparing replicated data drawn from the joint posterior predictive distribution to observed data. A measure of goodness-of-fit is the percentage of observed data falling within the 95% predicted credible intervals [23].

### WAIC and LOO-CV

Information criteria such as the WAIC and LOO-CV are common measures of predictive precision also used to compare models. The WAIC is the sum of the log predictive density computed for every point, to which a bias is added to take into account the number of parameters. The LOO-CV method uses the log predictive density estimated from a training subset and applies it to another one [23]. Both the WAIC and LOO-CV criteria were computed with the R package *bayesplot* [22].

### Mathematical definition and properties of *LC*(*x*, *t*)

The *LC*(*x*, *t*) makes sense only under conditions of constant exposure profiles (i.e., for any time *t*, *C*_*w*_(*t*) is constant). In such situations, we can provide an explicit formulation of the survival rate over time by considering both the GUTS-RED-SD and GUTS-RED-IT models. Many software provide an implementation of GUTS models that make it possible to compute the *LC*(*x*, *t*) at any time and for any *x*% [30]. Our Bayesian implementation of GUTS models using the R environment is one example [8].

Let *LC*(*x*, *t*) be the lethal concentration for *x*% of organisms at any time *t* and *S*(*C, t*) be the survival rate at the constant concentration *C* and time *t*. Then, the *LC*(*x*, *t*) is defined as

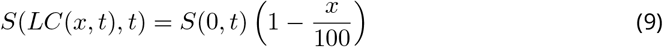

where *S*(0, *t*) is the survival rate at time *t* when there is no contaminant, which reflects the background mortality.

### GUTS-RED-SD model

The lethal concentration *LC*_*SD*_(*x*, *t*) is given by

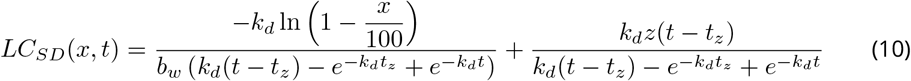

As mentioned in the Supplementary Material, under time-variable exposure, *t*_*z*_ also varies over time, while in the case of constant exposure, *t*_*z*_ is exactly −1/*k*_*d*_ ln(1 − *z*/*C*_*w*_). This expression of *t*_*z*_ prevents an explicit formulation of *LC*_*SD*_(*x*, *t*). For increasing time, the *LC*_*SD*_(*x*, *t*) curve becomes a vertical line at concentration *z*. We assume that the threshold concentration *z* is reached in a finite amount of time, which means that 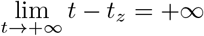.

Therefore, when time tends to infinity, the convergence is

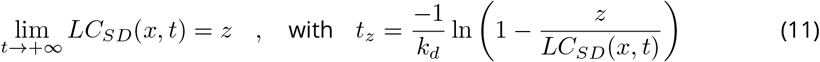

### GUTS-RED-IT model

The lethal concentration *LC_IT_* (*x*, *t*) is given by

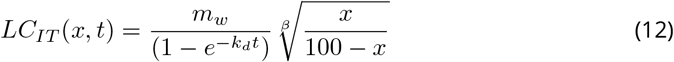

It is then clear that as *t* increases, the *LC*_*IT*_ (*x*, *t*) converges to

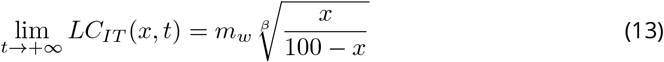

In the specific case of *x* = 50%, we get 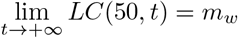.

### Calculation of the density distribution of *LC*(*x*, *t*)

The calculation of *LC*(*x*, *t*) is based on equation (9). Using the GUTS models and the estimates of parameters from the calibration processes, we compute the survival rate without contamination (i.e., the background mortality, denoted *S*(0, *t*)) and a set of predictions of the survival rate over a range of concentrations (i.e., *S*(*C*, *t*)).

### Mathematical definition and properties of the multiplication factor *MF* (*x*, *t*)

Contrary to the lethal concentration *LC*(*x*, *t*) used under conditions of constant exposure profiles, the multiplication factor *MF* (*x*, *t*) can be computed for both constant and time-variable exposure profiles.

With the exposure profile *C*_*w*_(*τ*), with *τ* ranging from 0 to *t*, the *MF* (*x*, *t*) is defined as

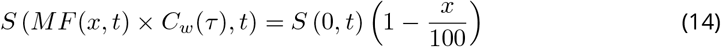

In the Supplementary Material, we show that the internal damage *D*_*w*_(*t*) is linearly related to the multiplication factor since regardless of the exposure profile (constant or time-variable), we get the following relationship:

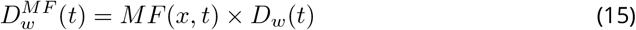

where 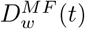 is the internal damage when the exposure profile is multiplied by *MF* (*x*, *t*).

### GUTS-RED-SD model

The multiplication factor *MF_SD_*(*x*, *t*) is given by

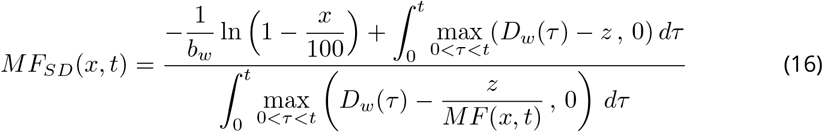

### GUTS-RED-IT model

The multiplication factor *MF_IT_* (*x*, *t*) is given by

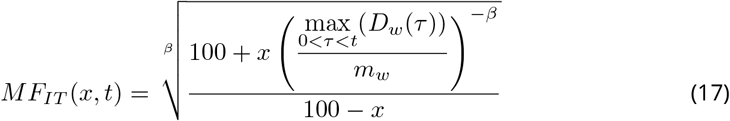

Therefore, from a GUTS-RED-IT model, solving the toxicokinetics part, which gives 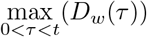, is enough to find any multiplication factor for any *x* at any *t*. When the external concentration is constant, this maximum is 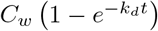.

## Results

### Goodness-of-fit of GUTS-RED-SD and GUTS-RED-IT models

For all compounds, fitting observed survival with test data obtained under constant exposure profiles provides better fits than using data from testing under time-variable exposure profiles (Table 2, see also posterior predictive check graphics in Supplementary Material), regardless of the measure of goodness-of-fit (except for the NRMSE measure used on the GUTS-RED-IT model of dimethoate). This result is unsurprising since, as shown in Table 1, there are always more time series in data sets with constant exposure profiles. However, since there are explicit solutions of differential equations with constant exposure profiles for both the GUTS-RED-SD and GUTS-RED-IT models, the computational process for constant exposure profiles is easier than that for time-variable exposure profiles, which requires the use of a numerical integrator.

For validation, we calibrated the model on a data set A to then predict another data set B. As a result, regardless of the measure of goodness-of-fit, the predictions are always better when the calibration is carried out using data of time-variable exposure profiles to then predict data from constant exposure profiles than when the inverse was carried out, that is, calibration using data from testing under constant exposure profiles to then predict data from testing under time-variable exposure profiles.

Table 2 shows that the GUTS-RED-SD and GUTS-RED-IT models are similar in the quality of their fits. However, the GUTS-RED-IT model particularly underperforms for carbendazim and dimethoate under time-variable exposure profiles. Nonetheless, under time-variable exposure profiles for the malathion and propiconazole data sets, the 95% credible interval for the GUTS-RED-IT model is large (see figures in the Supplementary Material). However, when uncertainties are large, the 95% credible interval around predictions used for the PPC tends to cover all the observations regardless of the fitting accuracy. The Bayesian measures WAIC and LOO-CV are better for penalizing excessively large uncertainties.

**Table 2.** Results of calibration and validation of the GUTS-RED-SD and GUTS-RED-IT models for the five chemical compounds: carbendazim (car), cypermethrin (cyp), dimethoate (dim), malathion (mal) and propiconazole (prz). Profiles of exposure concentrations are either constant, denoted *cst*, or variable, denoted *var*. The notation *cst* → *var* indicates that calibration was carried out with a data set of constant exposure and that validation was carried out with a data set of time-variable exposure (see data set in Table 1). The measures NRMSE, %PPC, WAIC and LOO-CV assess the goodness-of-fit and are fully explained in section.

| Product | Profile | GUTS SD |  |  |  | GUTS IT |  |  |  |
| --- | --- | --- | --- | --- | --- | --- | --- | --- | --- |
|  |  | NRMSE | %PPC | WAIC | LOO-CV | NRMSE | %PPC | WAIC | LOO-CV |
| Calibration |  |  |  |  |  |  |  |  |  |
| car | cst | 0.112 | 100 | 402.41 | 403.27 | 0.124 | 100 | 420.11 | 422.09 |
| cyp | cst | 0.095 | 100 | 196.37 | 206.78 | 0.092 | 100 | 188.07 | 189.09 |
| dim | cst | 0.122 | 97.5 | 308.94 | 309.41 | 0.171 | 90.0 | 357.38 | 358.74 |
| mal | cst | 0.090 | 100 | 248.87 | 249.59 | 0.112 | 92.5 | 273.01 | 273.54 |
| prz | cst | 0.102 | 100 | 282.03 | 285.57 | 0.118 | 80.0 | 308.03 | 314.93 |
| car | var | 0.159 | 82.1 | 1006.0 | 1012.1 | 0.499 | 32.1 | 1222.4 | 1216.4 |
| cyp | var | 0.196 | 91.7 | 829.04 | 833.48 | 0.116 | 97.2 | 793.95 | 801.23 |
| dim | var | 0.129 | 83.3 | 1224.8 | 1226.8 | 0.161 | 55.6 | 1357.2 | 1344.7 |
| mal | var | 0.196 | 97.8 | 762.58 | 766.76 | 0.148 | 100 | 908.56 | 934.80 |
| prz | var | 0.164 | 95.5 | 951.10 | 894.02 | 0.038 | 97.7 | 3262.8 | 1436.2 |
| Validation: data used for parameter calibration → data for prediction and goodness-of-fit |  |  |  |  |  |  |  |  |  |
| car | cst → var | 0.159 | 42.9 | 17709 | 4578.4 | 0.148 | 50.0 | 12800 | 4541.0 |
| cyp | cst → var | 0.196 | 91.7 | 1760.5 | 1423.5 | 0.183 | 88.9 | 1283.4 | 1141.0 |
| dim | cst → var | 0.129 | 83.3 | 1845.7 | 1685.3 | 0.199 | 63.9 | 1708.7 | 1628.9 |
| mal | cst → var | 0.196 | 67.4 | 10162 | 2610.7 | 0.169 | 63.0 | 1258.5 | 1286.1 |
| prz | cst → var | 0.164 | 95.5 | 940.54 | 900.90 | 0.176 | 90.9 | 894.41 | 940.74 |
| car | var → cst | 0.164 | 67.5 | 537.14 | 537.79 | 0.228 | 90.0 | 437.01 | 437.01 |
| cyp | var → cst | 0.071 | 82.5 | 537.62 | 488.90 | 0.051 | 87.5 | 453.65 | 378.89 |
| dim | var → cst | 0.013 | 97.5 | 302.24 | 302.30 | 0.157 | 87.5 | 389.32 | 393.68 |
| mal | var → cst | 0.053 | 80.0 | 470.28 | 512.86 | 0.049 | 90.0 | 869.45 | 732.94 |
| prz | var → cst | 0.040 | 77.5 | 797.60 | 660.09 | 0.041 | 80.0 | 1661.3 | 1107.8 |

### Comparison of *LC*(*x*, *t*) between GUTS-RED-SD and GUTS-RED-IT models

There is no obvious difference between the GUTS-RED-SD and GUTS-RED-IT models in their goodness-of-fit nor in the calculation of *LC*(*x*, *t*) over time *t* or for different percentages of the population affected (*x*).

**Figure 1.**
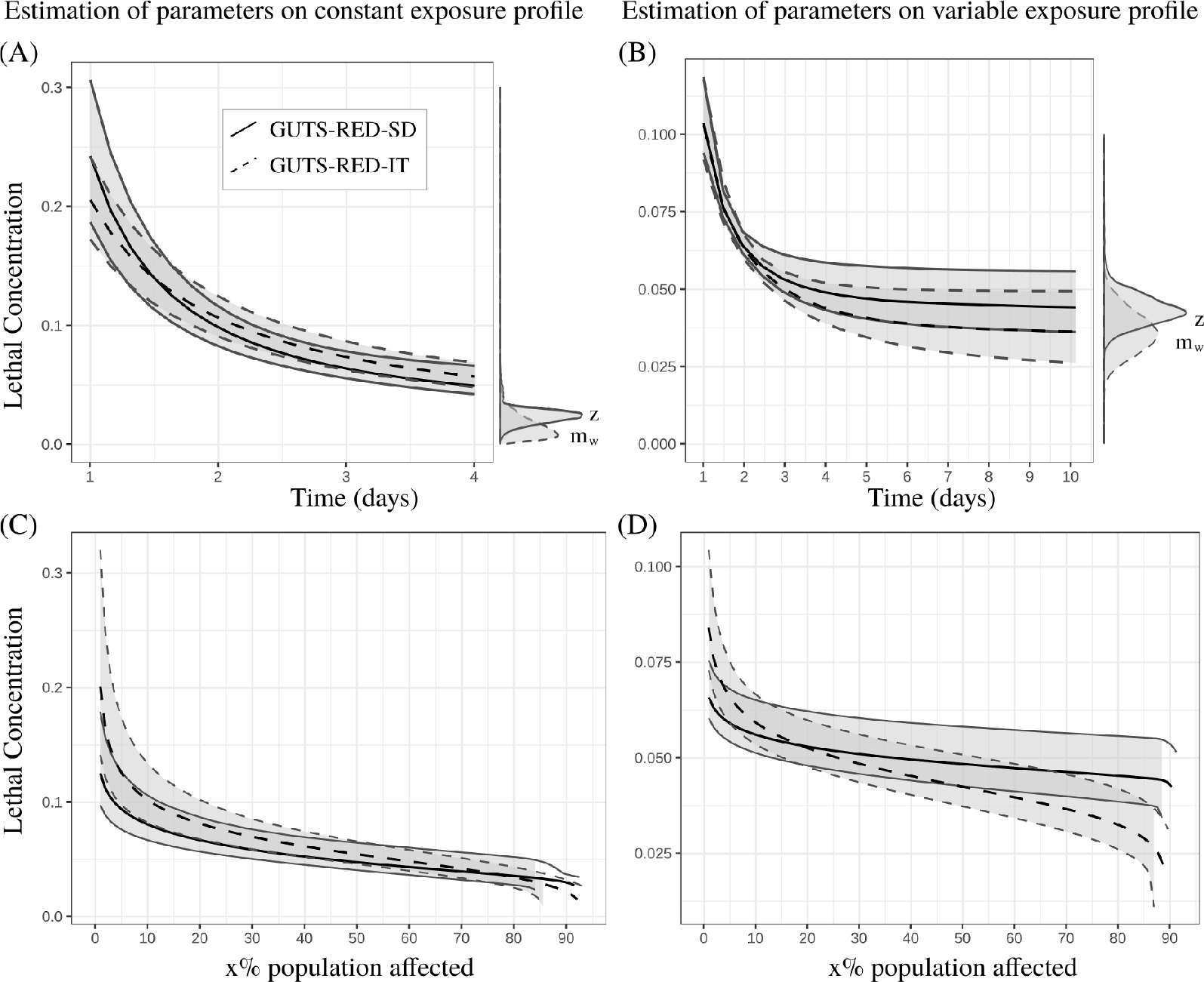
Comparison of *LC*(*x*, *t*) between GUTS-RED-SD, solid lines, and GUTS-RED-IT models, dashed lines, for cypermethrin (see Supplementary Material for other compounds). Parameters are estimated with data collected under constant (A, C) and variable (B, D) concentration profiles. Black lines are medians, and grey zones are 95% credible bands. (A, B) Lethal concentration for 50% of the organisms (*LC*(50, *t*)) from day 1 to the end of the experiment. (C, D) Lethal concentration at the end of experiment (4 and 10 days, respectively) against the percentage of the population affected.

### *LC*(*x*, *t*) as a function of time *t*

As expected, Figures 1-(A,B) and the Supplementary Material show that *LC*(*x*, *t*) decreases with time. The shape of this decrease, which is exponential and converges toward different values according to the model, is rarely analysed. This asymptotic behavior is known as the incipient *LC*(*x*, *t*) [31]. A direct consequence for risk assessors is that the evaluation of *LC*(*x*, *t*) at an early time induces higher sensitivity to time *t* than that at a later time (with the specific time being relative to the species and the compound). In other words, the sensitivity of *LC*(*x*, *t*) to time *t* decreases as long as *t* increases. For instance, Figures 1-(A,B) reveal that a small amount of change in time around day 2 leads to a greater change in the estimation of *LC*(*x*, *t*) than does a small amount around day 4. However, note that the uncertainty of *LC*(*x*, *t*) does not always decreases when time increases. For instance, as shown in Figure 1-(B), the uncertainty at day 6 and afterward is greater than that around day 3.

When *t* increases to infinity, *LC*(*x*, *t*) converges towards the distribution of parameter *z* for the GUTS-RED-SD model (see equation (11)) and 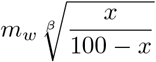 for the GUTS-RED-IT model (see equation (13)). The specific *LC*_50,*t*_ tends to *z* for the GUTS-RED-SD model and to *m*_*w*_ for the GUTS-RED-IT model (see equations (11) and (13)).

### *LC*(*x*, *t*) as a function of percentage of the population affected, *x*

As shown in Figure 1-(C,D), the uncertainty of *LC*(*x*, *t*) is greater at low values of *x*, that is, when the effect of the contaminant is weak. Although computing *LC*(*x*, *t*) at *x* > 50% is never used for ERA, the uncertainty of *LC*(*x*, *t*) increases when *x* tends to 100%. As a consequence, while the uncertainty is not always minimal at the standard value of *x* = 50%, it seems to always be smaller around this value than around *x* = 10%, another classical value used in ERA. Consequently, while TKTD models allow risk assessors to compute the *LC*(*x*, *t*) for any value of *x*, if only one value has to be chosen, we recommend that the standard *x* = 50% be chosen.

### Comparison of *MF* (*x*, *t*) between GUTS-RED-SD and GUTS-RED-IT models

#### *MF* (*x*, *t*) as a function of time *t*

As expected, Figures 2-(D-F) show that the multiplication factor decreases, or stay constant, when the time at which the survival rate is checked increases. In other words, the later the survival rate is assessed, the lower the multiplication factor is. In addition, these graphics reveal that there is no typical pattern in the curves of multiplication factors over time *t* of exposure. Under a constant exposure profile, the curve shows an exponential decreasing pattern, while under pulsed exposure, it shows a constant phase and, at the time when exposure peaks, a sudden decrease in the multiplication factor. The multiplication factor is clearly highly variable around a concentration pulse of the chemical product.

**Figure 2.**
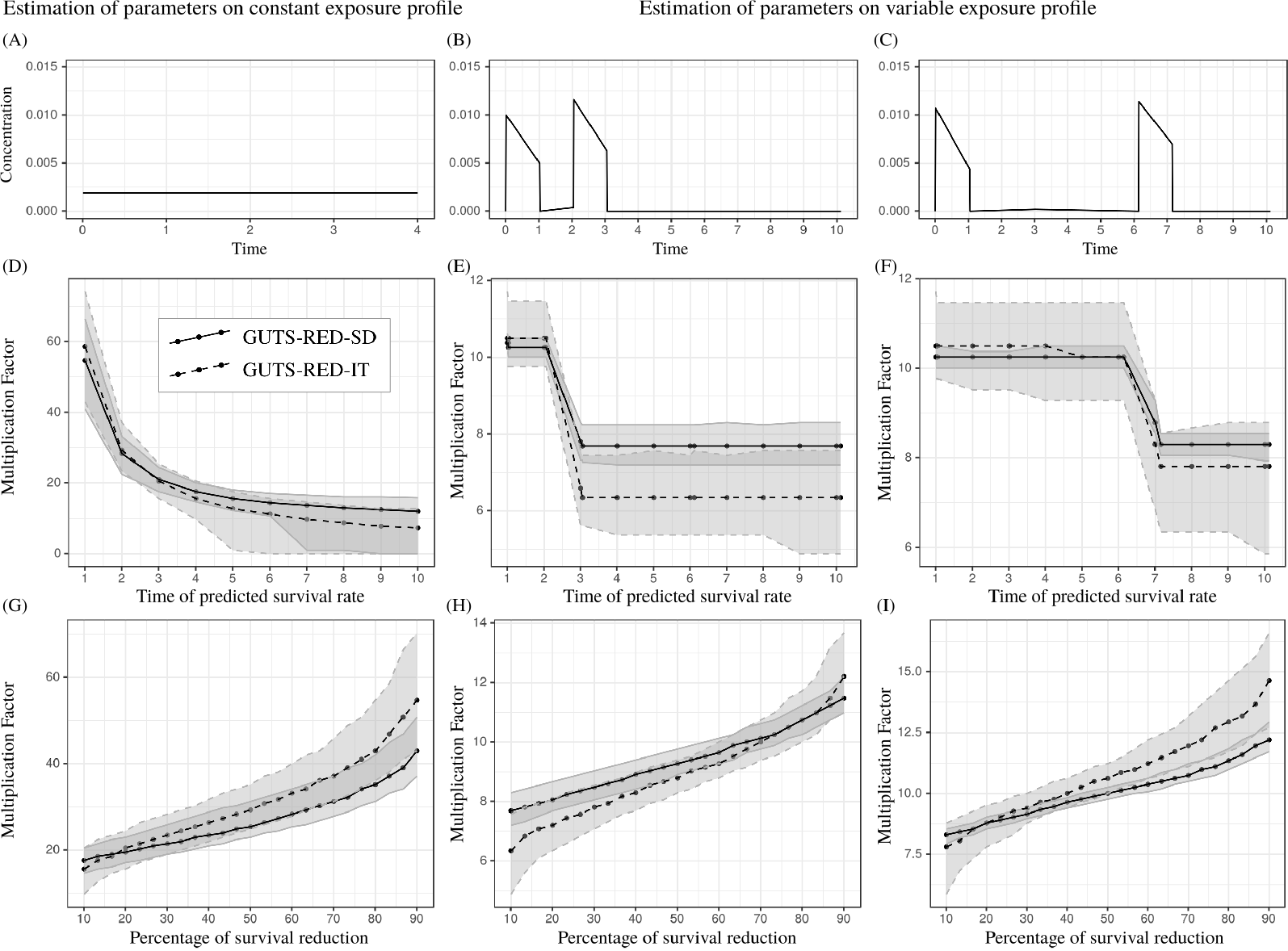
Comparison of *MF* (*x*, *t*) between GUTS-RED-SD, solid lines, and GUTS-RED-IT models, dashed lines, for cypermethrin (see Supplementary Material for other compounds). Parameters are estimated with data collected under constant (A, D, G) and variable (B, C, E, F, H, I) concentration profiles. (A-C) Exposure profiles, (D-F) Multiplication factors estimated for a 10% reduction in survival (i.e., *MF* (*x* = 10, *t*)) over time. (G-I) Multiplication factors estimated at the end of experiments (time=4 for (G) and 10 for (H, I)) against the percent survival reduction.

#### *MF* (*x*, *t*) as a function of percent survival reduction *x*

Unsurprisingly, Figures 2-(G-I) show that the multiplication factor increases with an increase in the percent reduction in the survival rate. An interesting result is the non-linearity of this increase. As observed for the *LC*(*x*, *t*), the uncertainty is greater at low and high percentages than for intermediate values near a 50% survival reduction. As a consequence, it would be relevant to set 50% as a standard for ERA.

**Figure 3.**
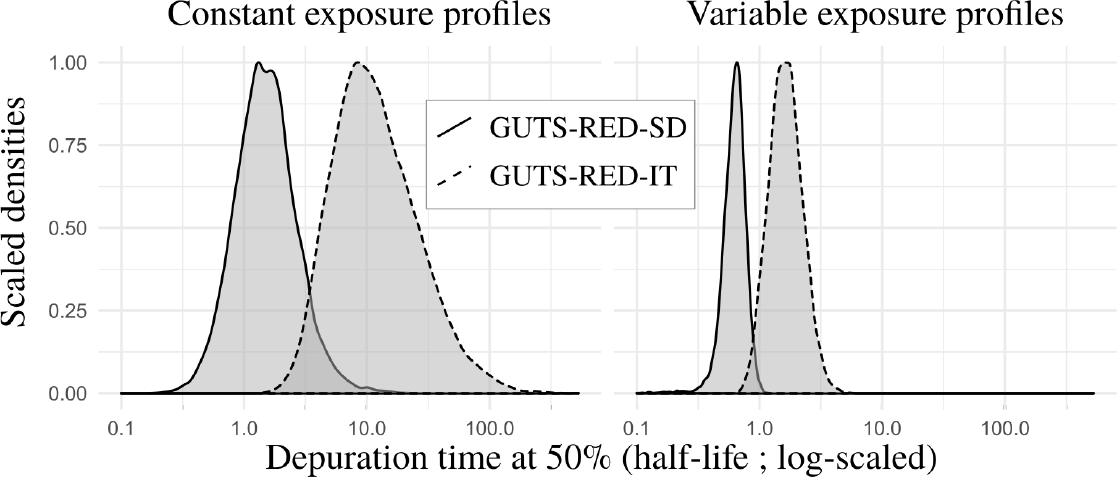
Distribution of estimated depuration time (see equation (3)) for cypermethrin in GUTS-RED-SD and GUTS-RED-IT models for data sets collected under constant (left) and variable (right) exposure profiles. Median and 95% credible interval of the 50% depuration time are 1.48 [0.502, 5.00] under constant exposure profiles for the GUTS-RED-SD model and 10.8 [3.21, 68.5] under constant exposure profiles for the GUTS-RED-IT model, and those under variable exposure profiles are 0.633 [0.386, 0.890] for the GUTS-RED-SD model and 1.62 [0.917, 3.06] for the GUTS-RED-IT model.

### Effect of the depuration time on the predicted survival rate

#### Patterns of internal scaled concentrations

The dominant rate constant *k*_*d*_, which regulates the kinetics of the toxicant, is always greater for the GUTS-RED-SD model than for the GUTS-RED-IT model, such that the depuration time for the GUTS-RED-SD model is always smaller than that for the GUTS-RED-IT model (see Figure 3 and Supplementary Material). As a consequence, under a time-variable exposure concentration, the internal scaled concentration with the GUTS-RED-SD model has a greater amplitude than that with the GUTS-RED-IT model (Figures 4, 5 and Supplementary Material). In other words, the toxicokinetics with the GUTS-RED-IT model are smoother than those with the GUTS-RED-SD model. Compensation for differences in *k*_*d*_ and therefore in the scaled internal concentrations comes from the other parameters: the threshold *z* and the mortality rate *k*_*k*_ for the GUTS-RED-SD model and the median threshold *m*_*w*_ and shape *β* for the GUTS-RED-IT model. However, when the calibration of the models is based on the same observed number of survivors, the threshold parameter *z* for the GUTS-RED-SD model and the median threshold *m*_*w*_ for the GUTS-RED-IT model are shifted.

#### Variation in the number of pulses in exposure profiles

The first step has been to explore the effect of the number of pulses (9, 6 and 3 pulses of one day each) over a period of 20 days with the same cumulative amount of contaminant in the external concentration after the 20 days (Figure 4 and Supplementary Material). For a conservative approach for ERA, regardless of whether the GUTS-RED-SD or GUTS-RED-IT model is used, it seems better to have few pulses of high amplitude than many pulses of low amplitude. Indeed, the survival rate over time with only 3 high pulses is lower than the survival rate under frequent lower exposure. This difference is confirmed in the Supplementary Material for the malathion and propiconazole data sets. Since the cumulative amount of contaminant is not changed, we do not see any effect of contaminant depuration (equation (3) and Figure 3), which could help individuals recover under a lower frequency of peaks. The comparison between constant and time-variable exposure profiles (Figure 4 and Supplementary Material) suggests that uncertainty is smaller when calibration is performed with data collected under a time-variable exposure profile. This result is counter-intuitive, especially since the number of time series was higher for the constant exposure profiles, which would reduce the uncertainties of parameter estimates. If this result is confirmed, then it would be better to predict variable exposure profiles with parameters calibrated from time-variable exposure data sets.

**Figure 4.**
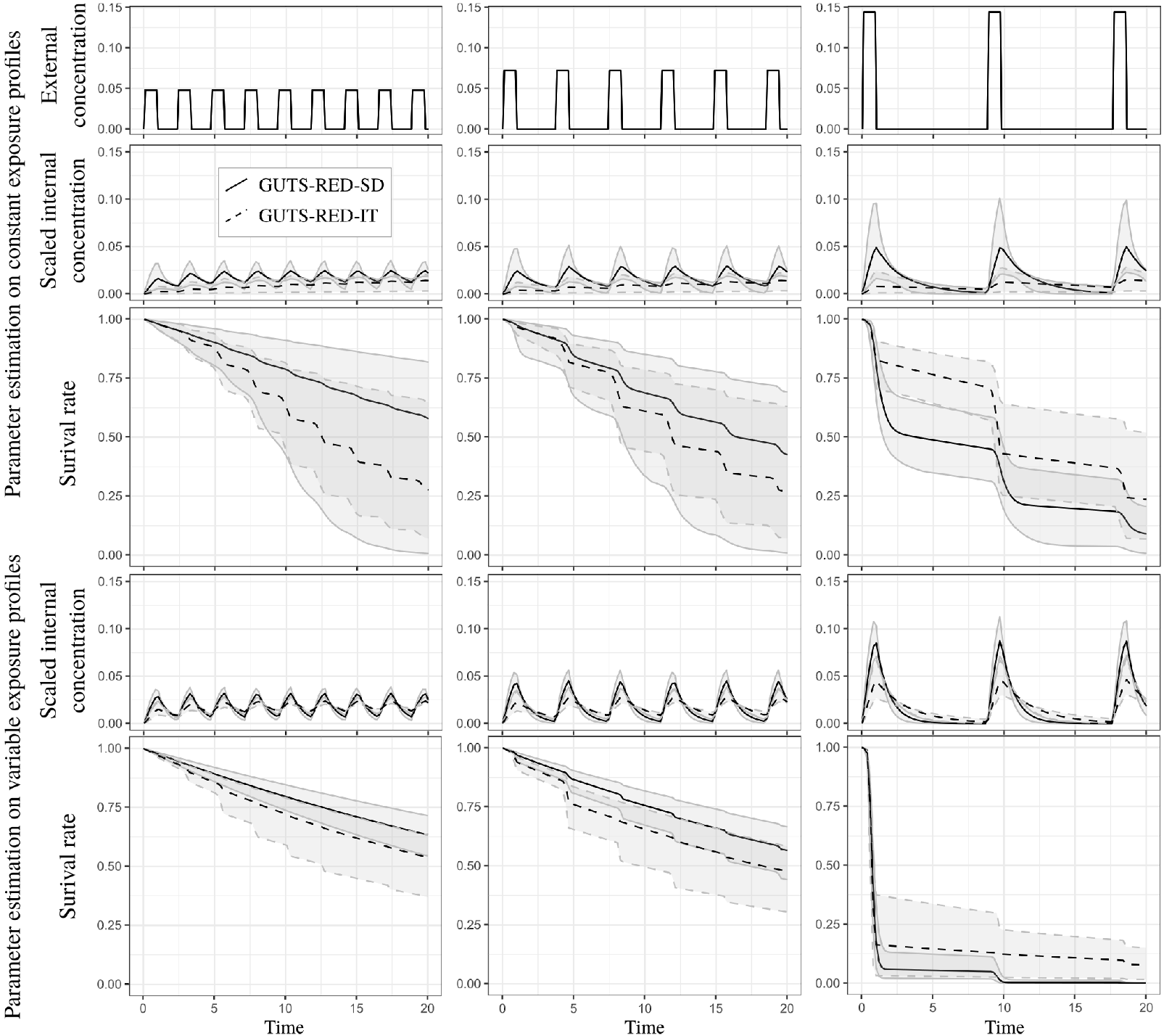
Survival rate over time with GUTS-RED-SD and GUTS-RED-IT models (solid and dashed lines, respectively) under different exposure profiles with the same area under the curve (with differences in the time after pulses and the maximal concentration of pulses). Parameters were estimated from the cypermethrin data set, under either constant (upper panel of the figure) or time-variable (lower panel of the figure) exposure.

**Figure 5.**
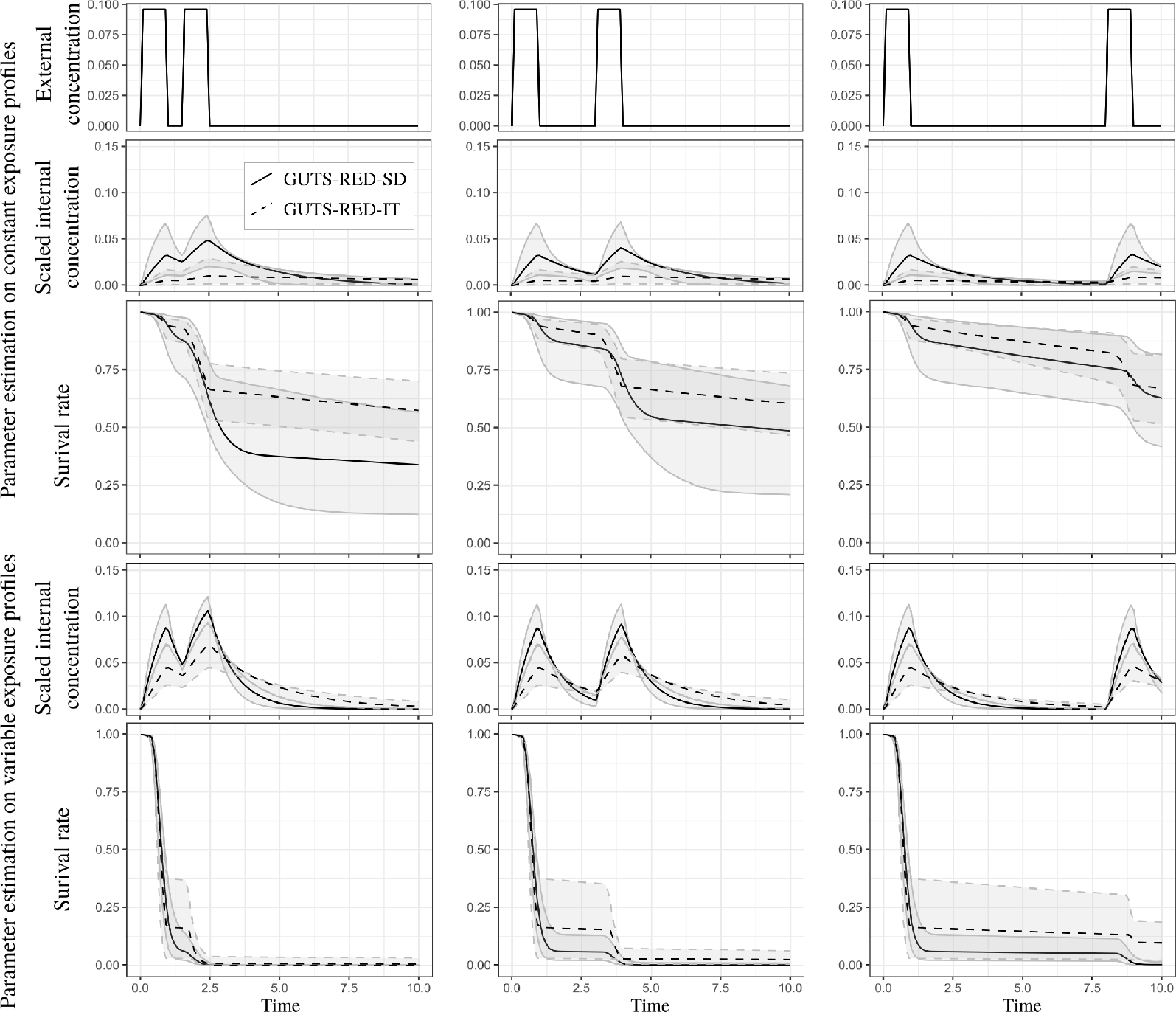
Survival rate over time with GUTS-RED-SD and GUTS-RED-IT models (solid and dashed lines, respectively) under a two-pulse exposure profile with the same area under the curve (with differences in the time between the two pulses). Parameters were estimated from the cypermethrin data set, under either constant (upper panel of the figure) or time-variable (lower panel of the figure) exposure.

#### Variation in the period between two pulses

To explore the effect of depuration time, we simulated exposure profiles under two pulses with different periods of time between them (i.e., 1/2, 2 or 7 days). The cumulative amount of contaminant remained the same for the three simulations. Figure 5 shows that increasing the period between two pulses may increase the survival rate of individuals, regardless of whether the GUTS-RED-SD or GUTS-RED-IT model is used. This is a typical result of extending the depuration period, which reduces the level of scaled internal concentration and therefore reduces the damage. We can easily see that the highest scaled internal concentration is reached when the pulse interval is the smallest. In this scenario, the addition of damages from the two pulses is clear. Again, because of the different depuration times of the two GUTS models, the results are different.

## Discussion

### Tracking uncertainties for environmental quality standards

Regardless of the scientific field, risk assessment is by definition linked to the notion of probability, characterized by different uncertainties such as the variability among organisms and noise in observations. In this sense, tracking how uncertainty propagates into models from collected data to model calculations of toxicity endpoints that are finally used for EQSs derivation is of fundamental interest for ERA [14]. For ERA, achieving good fits of experimental data is not enough. Instead, the key objective is the application of these fits to predict adverse effects under real environmental exposure profiles and to derive robust EQSs [19, 20, 24, 28, 32]. In this context, as we have shown in this paper, TKTD models calibrated under a Bayesian framework have two great advantages: on the one hand, TKTD models allow predictions of regulatory toxicity endpoints under any type of exposure profile; on the other hand, the Bayesian approach provides the marginal distribution of each parameter, and in this way, allows one to track the uncertainty of any prediction of interest.

Previous studies investigating goodness-of-fit did not find typical differences between GUTS-RED-SD and GUTS-RED-IT models [7, 10]. Our study confirms that under the specific consideration of uncertainties in regulatory toxicity endpoints, there is no evidence to support choosing either the GUTS-RED-SD or GUTS-RED-IT model over the other. A simple recommendation is therefore to use both and then, if they are successfully validated, take the most conservative scenario in terms of the ERA. With the 10 data sets we used and the 20 fittings we performed, the four measures of goodness-of-fit showed similar outputs for the GUTS-RED-SD and GUTS-RED-IT models under both constant and time-variable exposure profiles. The percentage of observed data falling within the 95% predicted credible interval, %*PPC*, has the advantage of being linked to visual graphics, i.e., PPC plots, and is therefore easier for risk assessors and stakeholders to interpret than the Bayesian WAIC and LOO-CV measures [12]. However, when the uncertainty is very large, predictions with their 95% credible intervals are likely to cover all of the observations, even in cases of low model accuracy. We showed that the WAIC and LOO-CV criteria are more robust probability measures for penalizing fits with large uncertainties [23]. Since the NRMSE is easy to calculate for any inference method (e.g., maximum likelihood estimation), it is also a relevant measure for checking the goodness-of-fit of models, as recently recommended by [20].

### What about the use and abuse of the lethal concentration?

After checking the quality of model parameter calibration, the next question is about the uncertainty of toxicity endpoints used to derive EQSs. Lethal concentrations are currently a standard for hazard characterization at the levels of a 10, 20 and 50% effect on the population. We show that the uncertainty of lethal concentrations differs according to the percentage *x* under consideration (Figure 1). It appears that this uncertainty is maximal at the extremes (toward 0 and 100%) and limited around 50%. Since the point of minimal uncertainty may drastically change depending on the experimental design, it could be relevant to extrapolate the lethal concentration for a continuous range of *x* (e.g., 10 to 50%), as we did for Figures 1-(C,D).

Many criticisms have targeted the lethal and effective concentrations for *x*% of the population and other related measures [28]. For instance, the classical way to compute the lethal concentration, at the final time point, ignores information provided by the observations made throughout the experiment and thus hides the time dependency. For the lethal effect, a classical approach to limit the variability in the period of time is to consider a long enough exposure duration to obtain the incipient lethal concentration (i.e., *LC*(*x*, *t* → +∞)) [31], that is, when the lethal concentration reaches its asymptote and no longer changes with an increasing duration of exposure, as observed in Figure 1. We provide mathematical expression of the lethal concentration convergence and explicit results when *x* = 50% for both GUTS models. We can therefore use the joint posterior parameter distribution provided by Bayesian inference to compute the distribution of the incipient lethal concentration.

A consequence of the exponential decrease in the lethal concentration with increasing time is that the sensitivity to time is greater early on, when a small change in time induces a great change in the lethal concentration regardless of *x*. Our analysis confirms that the classical evaluation of lethal concentration at the last time point of an experiment is supported by theoretical considerations. Hence, when comparing the lethal concentrations of different compounds or species that may require different experiment durations, using TKTD to extrapolate to other time points is highly advantageous.

### What does it mean to use a margin of safety?

Among the criticisms of the lethal concentration, one is that it is meaningful only under a set of constant environmental conditions, including a constant exposure profile [28, 31]. When the concentration of chemical compounds in the environment is highly variable over time, the use of toxicity endpoints based on toxicity data for constant exposure profiles may hide some processes, such as the response to pulses of exposure. This inadequacy is the reason underlying the interest in multiplication factors for ERA [7, 20].

A margin of safety deduced from a multiplication factor quantifies how far the exposure profile is below toxic concentrations [7]. Then, a key objective for risk assessors is to target the safest exposure duration and percentage effect on survival, *x*. Our study reveals a lower uncertainty around an *x* value of 50%. Thus, to reduce the uncertainty of the multiplication factor estimation, we recommend that 50% be selected, at least for comparisons between studies. We also show that under constant exposure profiles, the multiplication factor exhibits an asymptotic shape similar to that of the lethal concentration. There is an incipient value of the multiplication factor for any *x* as time goes to infinity. Therefore, under constant profiles, we recommend that the latest time point in the exposure profile be used to determine toxicity endpoints to reduce the sensitivity of the multiplication factor estimation to time.

However, the multiplication factor is meaningful when applied to realistic exposure profiles, which are rarely constant, and our study shows that there is no asymptotic shape under such conditions. In addition, we observed great sensitivity of the multiplication factor to time around peaks in the exposure profiles, that is, high variation in the multiplication factor with a small amount of change in time. Therefore, it is recommended that multiplication factors be computed only some time (e.g., several days) after a peak. More generally, the multiplication factor is designed to be compared to the assessment factor (AF) classically used with the effect/lethal concentration value to derive EQSs based on real-world exposure profiles. As a consequence, assessors must be very careful in examining the characteristics of pulses in the exposure profiles (e.g., frequencies and amplitudes) to understand how they drive changes in the multiplication factor. For such exploration, taking advantage of TKTD capabilities to generate predictions at any time is valuable.

### Effect of depuration in time-variable exposure profiles

The survival response to pulses depends on the depuration time driven by the toxicokinetics part of the TKTD model. The kinetics of assimilation and elimination of compounds integrated within the toxicokinetic module are a fundamental part of ecotoxicological models [38]. In reduced GUTS models, namely, GUTS-RED-SD and GUTS-RED-IT models, we assume no measure of the internal concentration, which is therefore calibrated at the same time as other parameters included in the toxicodynamics part. The resulting scaled damage is defined by the toxicodynamics, for which there are two different hypotheses regarding the mechanism of mortality for GUTS-RED-SD and GUTS-RED-IT models. As a consequence, our results illustrate that the scaled damage does not have the same meaning in GUTS-RED-SD and GUTS-RED-IT models and therefore cannot be directly compared between them.

In both models, from the underlying mechanism, we know that damage is positively correlated with pulse amplitude: the lower the amplitude is, the lower the damage is, as shown in Figure 4. As a result, for the same cumulative amount of contaminant in an experiment, using fewer pulses reduces final survival rates. Therefore, the most conservative experimental design is one with fewer pulses of relatively high amplitude.

Furthermore, in Figure 5, we bring to light the effect of depuration time. When pulses are close together, the organisms do not have time to depurate; therefore, the damage accumulates and thus has a cumulative effect on survival. As a consequence, in a long enough experiment, when pulses become less correlated in terms of cumulative damage, then the final survival rate increases. Because of this phenomenon, we recommend an experimental design with two close pulses, as it is the more conservative in terms of ERA. However, to achieve better calibration of the toxicokinetic parameter, which would potentially differentiate the GUTS-RED-SD model from the GUTS-RED-IT one, it is important to also include uncorrelated pulses in the experimental design.

Finally, our study reveals that the uncertainty of predictions under time-variable exposure profiles seems to be smaller when calibration is performed with data sets under time-variable rather than constant exposure profiles. While this observation makes theoretical sense, since predictions are made with the same type of profile as that used for calibration of the parameters, further empirical studies must be performed to confirm this point.

The environmental dynamics of chemical compounds can be highly variable depending not only on the whole environmental context (e.g., anthropogenic activities, geochemical kinetics, and ecosystem processes) but also on the chemical and biological transformation of the compound under study. Therefore, as a general recommendation, we would like to point out the relevancy of experimenting with several type of exposure profiles. Generally, a control and both constant and time-variable exposure profiles including toxicologically dependent and independent pulses seem to be the minimum requirements.

### Practical use of GUTS models

#### Optimization and exploration of experimental designs

The complexity of environmental systems combined with thousands of compounds produced by human activities implies the need to assess environmental risk for a very large set of species-compound combinations [6]. As a direct consequence, optimizing experimental design to maximize the gain in high-quality information from experiments is a challenging requisite for which mechanism-based models combined with a Bayesian approach offer several tools [1]. An extension of the present study would be to use the joint posterior distribution of parameters and the distribution of toxicity endpoints to quantify the gain in knowledge of several potential experiments to select the most informative. The next objective is thus to develop a framework that could help in the construction of new experimental designs to minimize their complexity and number while maximizing the robustness of toxicity endpoint estimates.

Despite their many advantages, TKTD models and therefore GUTS models remain little used. This lack of use is due to the mathematical complexity of such models based on differential equations that need to be numerically integrated when fitted to data [2]. By promoting GUTS models within regulatory documents associated with ERAs, the models could be further extended when available within a software environment allowing their implementation without the need to engage with technicalities. Currently, several software allow these difficulties to be circumvented [3, 8, 30], and a web platform has been proposed [11].

#### Limitations

Survival is the most often measured response to chemical toxins in the environment, but it may be more relevant to manage sub-lethal effects in ERA to prevent community collapse [9]. While the lethal concentration decreases as time increases, other sub-lethal effects (e.g., reproduction and growth) do not always follow this pattern [4, 28]. The concentration levels in acute toxicity tests are higher than those classically observed in the environment. Therefore, under real environmental conditions, sub-lethal effects may have more direct impacts on population dynamics than on survival. Thus, it would be of real interest to encompass different effects in a global TKTD approach to generate better predictions when scaling up to the population and community levels [28] and at multi-generationnal scales [14].

Another well-known limitation is the derivation of EQSs from specific species-compound combinations. To extrapolate ecotoxicological information from a set of single species tests to a community, ERA uses a species sensitivity (weighted) distribution (SS(W)D) which can be used to derive EQSs covering a set of taxonomically different species [16]. This calculation is classically applied to *LC*(*x*, *t*) and could easily be performed with *MF* (*x*, *t*) with the benefit of being applicable to time-variable exposure profiles [20].

## Conclusion

As recently written by EFSA experts, “*uncertainty analysis is the process of identifying limitations in scientific knowledge and evaluating their implications for scientific conclusions*” [18]. Inspired by the recent EFSA Scientific Opinion [20], we evaluated a combination of mechanism-based models with a Bayesian inference framework to track uncertainties of toxicity endpoints used in regulatory risk assessment with one compound-one species survival bioassays. We showed that the degree of uncertainty can change dramatically with time and depending on the exposure profile, revealing that single values such as the mean or median may be totally irrelevant for decision making. Description of uncertainties also increases transparency and trust in scientific outputs and is therefore key in applied sciences such as ecotoxicology. Many other kinds of uncertainties emerge along the decision chain, from the hazard identification to the characterization of risk. Focusing on uncertainty, such as through a Bayesian approach, should be a concern at every step and, above all, for any information returned by mathematical-computational models.

## Data accessibility

Data treatment and statistical scripts in R are fully available on the open data repository Github published though Zenodo – doi: https://zenodo.org/record/1972932

## Acknowledgements

The authors are very grateful for inputs from Theo Brock on an earlier version of the manuscript. We thank Andreas Focks and two anonymous reviewers for their valuable suggestions. The authors also thank the French National Agency for Water and Aquatic Environments (ONEMA, now the French Agency for Biodiversity) for its financial support. The authors declare no competing interests.

This preprint has been reviewed and recommended by Peer Community In Ecology – doi: https://dx.doi.org/10.24072/pci.ecology.100007

## Appendix

The two supplementary materials are available on BioRxiv – doi: https://doi.org/10.1101/356469

