## Supplementary material for "Recommendations to address uncertainties in environmental risk assessment using toxicokinetics-toxicodynamics models" (from another version of this preprint: version 2, 2018-11-22)

- Results of fit
  - Fitting plot
  - PPC: Posterior Predictive Check
  - Posteriors vs. Priors
  - Pairs plots
- LCx
  - LCx at final time
  - LCx along Time
  - LCx along X
- MFx
  - MFx along Time
  - MFx along X
- Predictions
  - Depuration times for every compounds
  - Prediction with different pulses frequencies and amplitudes

### Results of fit

#### Fitting plot

#### PPC: Posterior Predictive Check

#### Posteriors vs. Priors

#### Pairs plots

### LCx

#### LCx at final time

#### LCx along Time

#### LCx along X

### MFx

#### MFx along Time

#### MFx along X

### Predictions

#### Depuration times for every compounds

#### Prediction with different pulses frequencies and amplitudes
