## Supplementary material for "Recommendations to address uncertainties in environmental risk assessment using toxicokinetics-toxicodynamics models" (from another version of this preprint: version 2, 2018-11-22)

**Corresponding author** Sandrine Charles

### 1 Mathematical notations in reduced GUTS models

In this Supplementary Material concerning the derivation of mathematical equations, we use the following notations:

Table S1: Mathematical notations and descriptions

| Notation | Description |
|---|---|
| $h_b$ | Background hazard rate (intrinsic mortality rate) |
| $k_d$ | Dominant rate constant for the reduced GUTS model |
| $b_w$ | Killing rate, referenced to external concentration |
| $z$ | Threshold for effect |
| $m_w$ | Median of the distribution of threshold |
| $\beta$ | Shape parameter for the distribution threshold |
| $C_w$ | External chemical concentration in the environment |
| $D_w$ | Scaled damage, referenced to external concentration |
| $S(C_w, t)$ | Survival probability in a population of individuals ; $S_{SD}()$, $S_{IT}()$ with models $GUTS-RED-SD$, $GUTS-RED-IT$, reps. |
| $LC(x, t)$ | Lethal Concentration for $x\%$ of organisms at time $t$ ; $LC_{SD}()$, $LC_{IT}()$ with models $GUTS-RED-SD$, $GUTS-RED-IT$, reps. |
| $MF(x, t)$ | Multiplication Factor leading to an increase of $x\%$ of additional mortality at time $t$ ; $MF_{SD}()$, $MF_{IT}()$ with models $GUTS-RED-SD$, $GUTS-RED-IT$, reps. |

The dynamic of the internal damage (i.e. the toxicokinetics) is given by:

$$\frac{dD_w(t)}{dt} = k_d\left(C_w(t) - D_w(t)\right) \tag{S1}$$

We also suppose that $D_w(0) = 0$. Therefore, when the external concentration is constant (i.e. $C_w(t) = C_w$, we have:

$$D_w(t) = C_w \left(1 - e^{-k_d t}\right) \tag{S2}$$

Otherwise, if the external concentration is varying with time, we have:

$$D_w(t) = e^{-k_d t} k_d \int_0^t e^{k_d \tau} C_w(\tau)\, d\tau \tag{S3}$$

### 2 Mathematical analysis of $LC(x, t)$

The lethal concentration makes sense only in the case of constant exposure profile (i.e. $C_w(t) = C_w$, $\forall\, t$).

**Definition 1** ($LC(x,t)$)**.** *The lethal concentration for $x\%$ of organisms at time $t$ is the function denoted $LC(x,t)$ and defined (using notations provided in Table S1) as:*

$$S(LC(x,t), t) = S(0,t)\left(1 - \frac{x}{100}\right) \tag{S4}$$

*Where $S(LC(x,t),t)$ is the survival rate at time $t$ when the internal concentration is continuously the constant $LC(x,t)$ from time $0$ to time $t$. Also, $S(0,t)$ is the survival rate at time $t$ when there is no contaminant (i.e. external concentration equals $0$ all along the experiment), which is defined from the background mortality rate:*

$$S(0,t) = \exp\left(-\int_0^t h_b d\tau\right) = e^{-h_b t} \tag{S5}$$

#### 2.1 $LC_{SD}(x, t)$ for model GUTS-RED-SD

For GUTS-RED-SD model (with $S_{SD}(C,t)$ the survival rate for GUTS-RED-SD), the $LC_{SD}(x,t)$ is given by:

$$\begin{aligned}
& S_{SD}(LC_{SD}(x,t), t) = S_{SD}(0,t)\left(1 - \frac{x}{100}\right) \\
\Leftrightarrow\quad & \exp\left(-\int_0^t b_w \max_{0<\tau<t}(D_w(\tau) - z, 0) d\tau\right) e^{-h_b t} = e^{-h_b t}\left(1 - \frac{x}{100}\right) \\
\Leftrightarrow\quad & \int_0^t \max_{0<\tau<t}(D_w(\tau) - z, 0)\, d\tau = \frac{-1}{b_w} \ln\left(1 - \frac{x}{100}\right)
\end{aligned} \tag{S6}$$

Then, with $t_z$, the time at which the internal concentration is equal to the threshold, $D_w(t_z) = z$, using (S1), we have:

$$t_z = \frac{-1}{k_d} \ln\left(1 - \frac{z}{LC_{SD}(x,t)}\right) \tag{S7}$$

And equation (S6) can be developped as:

$$
\begin{aligned}
\text{(S6)} \Leftrightarrow \quad & \int_{t_z}^{t} \left(LC_{SD}(x,t)\left(1-e^{-k_d\tau}\right)-z\right)\,d\tau = \frac{-1}{b_w}\ln\left(1-\frac{x}{100}\right)\\
\Leftrightarrow \quad & LC_{SD}(x,t)(t-t_z) - z(t-t_z) + \frac{LC_{SD}(x,t)}{k_d}\left(e^{-k_dt}-e^{-k_dt_z}\right) = \frac{-1}{b_w}\ln\left(1-\frac{x}{100}\right)\\
\Leftrightarrow \quad & LC_{SD}(x,t)\left(t-t_z+\frac{1}{k_d}\left(e^{-k_dt}-e^{-k_dt_z}\right)\right) - z(t-t_z) = \frac{-1}{b_w}\ln\left(1-\frac{x}{100}\right)\\
\Leftrightarrow \quad & LC_{SD}(x,t) = \frac{-k_d\ln\left(1-\dfrac{x}{100}\right)}{b_w\left(k_d(t-t_z)-e^{-k_dt_z}+e^{-k_dt}\right)} + \frac{k_dz(t-t_z)}{k_d(t-t_z)-e^{-k_dt_z}+e^{-k_dt}}
\end{aligned}
\tag{S8}
$$

The expression of $t_z$ prevents to have an explicit formulation of $LC_{SD}(x,t)$.

We can use equation (S7) to see that:

$$
e^{-k_dt_z} = \exp\left(-k_d\frac{-1}{k_d}\ln\left(1-\frac{z}{LC_{SD}(x,t)}\right)\right) = 1-\frac{z}{LC_{SD}(x,t)} \tag{S9}
$$

Combining this result with equation (S8), we obtain:

$$
LC_{SD}(x,t) = \frac{\dfrac{-k_d}{b_w}\ln\left(1-\dfrac{x}{100}\right)+z(k_d(t-t_z)-1)}{k_d(t-t_z)+e^{-k_dt}-1} \tag{S10}
$$

##### 2.1.1 Convergence of $LC_{SD}(x,t)$ when $t$ tends to infinity

We assume the threshold concentration, $z$, is reached in a finite time, which mean that $\lim_{t\to+\infty} t-t_z=+\infty$.

To compute $\lim_{t\to+\infty} LC_{SD}(x,t)$, we have for one part of the equation:

$$
\lim_{t\to+\infty}\frac{\dfrac{-k_d}{b_w}\ln\left(1-\dfrac{x}{100}\right)}{k_d(t-t_z)+e^{-k_dt}-1} = 0 \tag{S11}
$$

And for the other part of the equation, we have:

$$
\lim_{t\to+\infty}\frac{z(k_d(t-t_z)-1)}{k_d(t-t_z)+e^{-k_dt}-1} = \lim_{t\to+\infty}\frac{z}{\dfrac{e^{-k_dt}}{k_d(t-t_z)-1}+1} = z \tag{S12}
$$

As a consequence, we obtain:

$$
\lim_{t\to+\infty} LC_{SD}(x,t) = z \tag{S13}
$$

##### 2.1.2 Influence of $x$ on the $LC_{SD}(x,t)$

**Sensitivity:**

To study the influence of $x$ on the $LC_{SD}(x,t)$, we compute the sensitivity, which is simply the derivative of $LC_{SD}(x,t)$ by $x$.

$$
\frac{\partial LC_{SD}(x,t)}{\partial x} = \frac{k_d}{(100-x)b_w\left(k_d(t-t_z)-e^{-k_dt_z}+e^{-k_dt}\right)} \tag{S14}
$$

Since $t \geq t_z$, then $b_w k_d(t-t_z) \geq 0$ while $e^{-k_d t} - e^{-k_d t_z} \leq 0$, and therefore the sign of the derivative may change according to the parameter values.

**Elasticity:**

The elasticity of $LC_{SD}$ to parameter $x$ represents the proportional change in $LC_{SD}$ in response to a proportional change in the parameter $x$ and is given by:

$$\frac{\partial LC_{SD}(x,t)}{\partial x} \times \frac{x}{LC_{SD}(x,t)} = \frac{x}{(100-x)\left(b_w z(t-t_z) - \ln\left(1-\frac{x}{100}\right)\right)} \tag{S15}$$

#### 2.2 $LC_{IT}(x,t)$ for model GUTS-RED-IT

For GUTS-RED-IT model (with $S_{IT}(C,t)$ the survival rate for GUTS-RED-IT), the $LC_{IT}(x,t)$ is given by:

$$\begin{aligned}
& S_{IT}(LC_{IT}(x,t),t) &&= S_{IT}(0,t)\left(1-\frac{x}{100}\right) \\
\Leftrightarrow\quad & e^{-h_b t}\left(1-\frac{1}{1+\left(\frac{LC_{IT}(x,t)(1-e^{-k_d t})}{m_w}\right)^{-\beta}}\right) &&= e^{-h_b t}\left(1-\frac{x}{100}\right) \\
\Leftrightarrow\quad & 1-\frac{1}{1+\left(\frac{LC_{IT}(x,t)(1-e^{-k_d t})}{m_w}\right)^{-\beta}} &&= 1-\frac{x}{100} \\
\Leftrightarrow\quad & 1+\left(\frac{LC_{IT}(x,t)(1-e^{-k_d t})}{m_w}\right)^{-\beta} &&= \frac{100}{x} \\
\Leftrightarrow\quad & \left(\frac{LC_{IT}(x,t)(1-e^{-k_d t})}{m_w}\right)^{-\beta} &&= \frac{100-x}{x} \\
\Leftrightarrow\quad & \left(\frac{LC_{IT}(x,t)(1-e^{-k_d t})}{m_w}\right)^{\beta} &&= \frac{x}{100-x} \\
\Leftrightarrow\quad & \frac{LC_{IT}(x,t)(1-e^{-k_d t})}{m_w} &&= \sqrt[\beta]{\frac{x}{100-x}} \\
\Leftrightarrow\quad & LC_{IT}(x,t) &&= \frac{m_w}{(1-e^{-k_d t})}\sqrt[\beta]{\frac{x}{100-x}}
\end{aligned} \tag{S16}$$

##### 2.2.1 Convergence of $LC_{IT}(x,t)$ when $t$ tends to infinity

Using equation (S16), we can direclty see that

$$\lim_{t\to+\infty} LC_{IT}(x,t) = m_w \sqrt[\beta]{\frac{x}{100-x}} \tag{S17}$$

For the specific case of $x = 50\%$, we have:

$$\lim_{t\to+\infty} LC_{IT}(50,t) = m_w \tag{S18}$$

###### 2.2.2 Influence of $x$ on $LC_{IT}(x,t)$

**Sensitivity:**
As for the $LC_{SD}(x,t)$, we compute the sensitivity of $LC_{IT}(x,t)$ for parameter $x$ by mean of the derivative:

$$\frac{\partial LC_{IT}(x,t)}{\partial x} = \frac{-100 m_w e^{k_d t} \sqrt[\beta]{\dfrac{x}{100-x}}}{\beta(100-x)x(1-e^{k_d t})} \tag{S19}$$

**Elasticity:**
Here, for this model, we can simplify the equation of sensitivity by computing the elasticity which is given by:

$$\frac{\partial LC_{IT}(x,t)}{\partial x} \times \frac{x}{LC_{IT}(x,t)} = \frac{-100 e^{k_d t}(1-e^{-k_d t})}{\beta(100-x)(1-e^{k_d t})} = \frac{100}{\beta(100-x)} \tag{S20}$$

The elasticity of $LC_{IT}(x,t)$ to parameter $x$ represents the proportional change of $LC_{IT}(x,t)$ in response to a proportional change in $x$. Here, the elasticity of $LC_{IT}(x,t)$ to $x$ is only function of $\beta$ and $x$.

### 3 Mathematical analysis of $MF(x,t)$

Contrary to the lethal concentration $LC(x,t)$, the multiplication factor, denoted $MF(x,t)$ makes sense for any type of exposure profile (constant or time-variable).

**Definition 2** ($MF(x,t)$)**.** *The multiplication factor leading to a reduction of $x\%$ of the survival rate at time $t$ is the function denoted $MF(x,t)$ and defined (using notations provided in Table S1) as:*

$$S\left(MF(x,t) \times C_w(\tau), t\right) = S\left(C_w(\tau), t\right)\left(1 - \frac{x}{100}\right) \tag{S21}$$

*where $C_w(\tau)$ is the exposure profile along the continuous period from 0 to $t$. The expression $S(MF(x,t) \times C_w(\tau), t)$ is the survival rate at time $t$ when the internal concentration is the exposure profile $C_w(\tau)$ multiplied by the constant multiplication factor $MF(x,t)$.*

#### 3.1 Internal concentration scaled by Multiplication Factor

The internal damage (or scaled internal concentration) at time $t$, $D_w(t)$, is given by equation (S1) using the profile of external concentration, $C_w(\tau)$, with $\tau$ going from 0 to $t$.

If the profile of external concentration is multiplied by a factor $MF(x,t)$ (assuming it reduces the survival rate by $x\%$ at time $t$), then, the internal concentration, denoted $D_w^{MF}(t)$, is also multiplied by this factor. From equation (S1) we get:

$$\frac{d D_w^{MF}(t)}{dt} = k_d(MF(x,t) \times C_w(\tau) - D_w^{MF}(t)) \quad \Rightarrow \quad D_w^{MF}(t) = MF(x,t) \times e^{-k_d t}\int_0^t k_d e^{k_d \tau} C_w(\tau) d\tau \tag{S22}$$

what leads to:

$$D_w^{MF}(t) = MF(x,t) \times D_w(t) \tag{S23}$$

Therefore, the internal damage referenced to the external concentration $D_w(t)$ is linearly scaled by the multiplication factor $MF(x,t)$ whatever the exposure profile $C_w(\tau)$.

#### 3.2 $MF_{SD}(x,t)$ for model GUTS-RED-SD

Using the definition of the multiplication factor, we have the following formulation for the GUTS-RED-SD model:

$$\begin{aligned}
& S_{SD}\left(MF_{SD}(x,t) \times C_w(\tau), t\right) = S_{SD}(C_w(\tau), t)\left(1 - \frac{x}{100}\right) \\
\Leftrightarrow\quad & \exp\left(-\int_0^t b_w \max_{0<\tau<t}(MF_{SD}(x,t)D_w(\tau) - z\,,\,0) + h_b\, d\tau\right) = \exp\left(-\int_0^t b_w \max_{0<\tau<t}(D_w(\tau) - z\,,\,0) + h_b\, d\tau\right)\left(1 - \frac{x}{100}\right) \\
\Leftrightarrow\quad & \int_0^t b_w \max_{0<\tau<t}(MF_{SD}(x,t)D_w(\tau) - z\,,\,0)\, d\tau = \int_0^t b_w \max_{0<\tau<t}(D_w(\tau) - z\,,\,0)\, d\tau - \ln\left(1 - \frac{x}{100}\right) \\
\Leftrightarrow\quad & b_w MF_{SD}(x,t)\int_0^t \max_{0<\tau<t}\left(D_w(\tau) - \frac{z}{MF_{SD}(x,t)}\,,\,0\right) d\tau = b_w \int_0^t \max_{0<\tau<t}(D_w(\tau) - z\,,\,0)\, d\tau - \ln\left(1 - \frac{x}{100}\right)
\end{aligned} \tag{S24}$$

When the external concentration is constant ($C_w(\tau) = C_w$), we can obtain $MF_{SD}(x,t)$ by using the explicit expression of $D_w(t)$ given by (S2). And so, we can define $t_z$ and $t_{z,MF}$ such as:

$$\begin{aligned}
& z = C_w\left(1 - e^{-k_d t_z}\right) && \Rightarrow \quad t_z = \frac{-1}{k_d}\ln\left(1 - \frac{z}{C_w}\right) \\
& \frac{z}{MF_{SD}(x,t)} = C_w\left(1 - e^{-k_d t_{z,MF}}\right) && \Rightarrow \quad t_{z,MF} = \frac{-1}{k_d}\ln\left(1 - \frac{z}{MF_{SD}(x,t)C_w}\right)
\end{aligned} \tag{S25}$$

Hence, we have:

$$\begin{aligned}
& b_w MF_{SD}(x,t)\int_0^t \max_{0<\tau<t}\left(D_w(\tau) - \frac{z}{MF_{SD}(x,t)}\,,\,0\right) d\tau = b_w \int_0^t \max_{0<\tau<t}(D_w(\tau) - z\,,\,0)\, d\tau - \ln\left(1 - \frac{x}{100}\right) \\
\Leftrightarrow\quad & b_w MF_{SD}(x,t)\int_{t_{z,MF}}^t C_w\left(1 - e^{-k_d\tau}\right) - \frac{z}{MF_{SD}(x,t)}\, d\tau = -\ln\left(1 - \frac{x}{100}\right) + b_w\int_{t_z}^t C_w\left(1 - e^{-k_d\tau}\right) - z\, d\tau \\
\Leftrightarrow\quad & b_w MF_{SD}(x,t)\left(\frac{C_w}{k_d}\left(e^{-k_d t} - e^{-k_d t_{z,MF}}\right) + \left(C_w - \frac{z}{MF_{SD}(x,t)}\right)(t - t_{z,MF})\right) = \\
& -\ln\left(1 - \frac{x}{100}\right) + b_w\left(\frac{C_w}{k_d}\left(e^{-k_d t} - e^{-k_d t_z}\right) + (C_w - z)(t - t_z)\right) \\
\Leftrightarrow\quad & MF_{SD}(x,t) = \frac{-\ln\left(1 - \frac{x}{100}\right) + b_w\left(\frac{C_w}{k_d}\left(e^{-k_d t} - e^{-k_d t_z}\right) + (C_w - z)(t - t_z)\right)}{b_w\left(\frac{C_w}{k_d}\left(e^{-k_d t} - e^{-k_d t_{z,MF}}\right) + \left(C_w - \frac{z}{MF_{SD}(x,t)}\right)(t - t_{z,MF})\right)}
\end{aligned} \tag{S26}$$

As for the $LC_{SD}(x,t)$, the expression of $t_{z,MF}$ prevents to have an explicit formulation of $MF_{SD}(x,t)$.

#### 3.3 $MF_{IT}(x,t)$ for model GUTS-RED-IT

Using the definition of the multiplication factor, we have the following formulation for the GUTS-RED-IT model:

$$
\begin{aligned}
&S_{IT}\left(MF_{IT}(x,t)\times C_w(\tau),t\right) = S_{IT}(C_w(\tau),t)\left(1-\frac{x}{100}\right)\\
\Leftrightarrow\quad & e^{-h_b t}\left(1-\frac{1}{1+\left(\frac{\max\limits_{0<\tau<t}\left(MF_{IT}(x,t)\times D_w(\tau)\right)}{m_w}\right)^{-\beta}}\right) = e^{-h_b t}\left(1-\frac{1}{1+\left(\frac{\max\limits_{0<\tau<t}\left(D_w(\tau)\right)}{m_w}\right)^{-\beta}}\right)\left(1-\frac{x}{100}\right)\\
\Leftrightarrow\quad & 1-\frac{1}{1+MF_{IT}(x,t)^{-\beta}\left(\frac{\max\limits_{0<\tau<t}\left(D_w(\tau)\right)}{m_w}\right)^{-\beta}} = \left(1-\frac{1}{1+\left(\frac{\max\limits_{0<\tau<t}\left(D_w(\tau)\right)}{m_w}\right)^{-\beta}}\right)\left(\frac{100-x}{100}\right)\\
\Leftrightarrow\quad & 1-\frac{1}{1+MF_{IT}(x,t)^{-\beta}\left(\frac{\max\limits_{0<\tau<t}\left(D_w(\tau)\right)}{m_w}\right)^{-\beta}} = \left(\left(\frac{\max\limits_{0<\tau<t}\left(D_w(\tau)\right)}{m_w}\right)^{-\beta}\frac{1}{1+\left(\frac{\max\limits_{0<\tau<t}\left(D_w(\tau)\right)}{m_w}\right)^{-\beta}}\right)\left(\frac{100-x}{100}\right)\\
\Leftrightarrow\quad & 1-\frac{1}{1+MF_{IT}(x,t)^{-\beta}\left(\frac{\max\limits_{0<\tau<t}\left(D_w(\tau)\right)}{m_w}\right)^{-\beta}} = \left(\frac{\max\limits_{0<\tau<t}\left(D_w(\tau)\right)}{m_w}\right)^{-\beta}\frac{(100-x)}{100+100\left(\frac{\max\limits_{0<\tau<t}\left(D_w(\tau)\right)}{m_w}\right)^{-\beta}}\\
\Leftrightarrow\quad & 1-\frac{1}{1+MF_{IT}(x,t)^{-\beta}\left(\frac{\max\limits_{0<\tau<t}\left(D_w(\tau)\right)}{m_w}\right)^{-\beta}} = \frac{100-x}{100+100\left(\frac{\max\limits_{0<\tau<t}\left(D_w(\tau)\right)}{m_w}\right)^{-\beta}}\\
\Leftrightarrow\quad & \frac{1}{1+MF_{IT}(x,t)^{-\beta}\left(\frac{\max\limits_{0<\tau<t}\left(D_w(\tau)\right)}{m_w}\right)^{-\beta}} = 1-\frac{100-x}{100+100\left(\frac{\max\limits_{0<\tau<t}\left(D_w(\tau)\right)}{m_w}\right)^{\beta}}\\
\Leftrightarrow\quad & 1+MF_{IT}(x,t)^{-\beta}\left(\frac{\max\limits_{0<\tau<t}\left(D_w(\tau)\right)}{m_w}\right)^{-\beta} = \frac{100+100\left(\frac{\max\limits_{0<\tau<t}\left(D_w(\tau)\right)}{m_w}\right)^{\beta}}{100+100\left(\frac{\max\limits_{0<\tau<t}\left(D_w(\tau)\right)}{m_w}\right)^{\beta}-(100-x)}\\
\Leftrightarrow\quad & MF_{IT}(x,t)^{-\beta}\left(\frac{\max\limits_{0<\tau<t}\left(D_w(\tau)\right)}{m_w}\right)^{-\beta} = \frac{100-x}{100+100\left(\frac{\max\limits_{0<\tau<t}\left(D_w(\tau)\right)}{m_w}\right)^{\beta}-(100-x)}\\
\Leftrightarrow\quad & MF_{IT}(x,t)^{-\beta} = \left(\frac{\max\limits_{0<\tau<t}\left(D_w(\tau)\right)}{m_w}\right)^{\beta}\frac{100-x}{100+100\left(\frac{\max\limits_{0<\tau<t}\left(D_w(\tau)\right)}{m_w}\right)^{\beta}-(100-x)}\\
\Leftrightarrow\quad & MF_{IT}(x,t)^{-\beta} = \frac{100-x}{100+x\left(\frac{\max\limits_{0<\tau<t}\left(D_w(\tau)\right)}{m_w}\right)^{-\beta}}\\
\Leftrightarrow\quad & MF_{IT}(x,t) = \sqrt[\beta]{\frac{100+x\left(\frac{\max\limits_{0<\tau<t}\left(D_w(\tau)\right)}{m_w}\right)^{-\beta}}{100-x}}
\end{aligned}
\tag{S27}
$$

Therefore, from a GUTS-RED-IT model, solving the toxicokinetics part giving $\max_{0<\tau<t}(D_w(\tau))$ is enough to find any multiplication factor for any $x$ at any $t$.

When the external concentration is constant (i.e., $C_w(\tau) = C_w$) we get:

$$MF_{IT}(x,t) = \left( \frac{100 + x \left( \frac{C_w \left(1 - e^{-k_d t}\right)}{m_w} \right)^{-\beta}}{100 - x} \right)^{1/\beta} \tag{S28}$$

### 4 Definition of WAIC and LOO-CV

For more information about information criteria and cross-validation, see [**?**]. Here we give a brief description to compute the Widely Available/Applicable Information Criterion (WAIC) and the Leave-One-Out Cross-Validation (LOO-CV).

#### 4.1 WAIC: Widely Available/Applicable Information Criterion

The first step is to compute the log pointwise predictive density:

$$lpd = \log \prod_{i=1}^{n} p_{post}(y_i) \Rightarrow \widehat{lpd} = \sum_{i=1}^{n} \log \left( \frac{1}{S} \sum_{s=1}^{S} p(y_i|\theta^S) \right) \tag{S29}$$

Then the variance of the log pointwise predictive density:

$$p_{WAIC} = \sum_{i=1}^{n} \mathbb{V}_{s=1}^{S}(\log p(y_i|\theta^s)) \tag{S30}$$

To finally compute the WAIC as:

$$WAIC = -2(\widehat{lpd} - p_{WAIC}) \tag{S31}$$

#### 4.2 LOO-CV: Leave-One-Out Cross-Validation

Cross-validation is like WAIC, applied several time on a subsample (S-1) drawn from the posterior.

$$\widehat{lpd_{-i}} = \sum_{j=1}^{n} \log \left( \frac{1}{S} \sum_{s=1}^{S} p(y_j|y_{-i};\theta^{is}) \right) \tag{S32}$$

Then the first order bias correction

$$\overline{lpd_{-i}} = \frac{1}{n} \sum_{i=1}^{n} \sum_{j=1}^{n} \log \left( \frac{1}{S} \sum_{s=1}^{S} p(y_j|y_{-i};\theta^{is}) \right) \tag{S33}$$

To finally compute the LOO-CV as:

$$LOO = -2 \left( \widehat{lpd_{-i}} + \widehat{lpd} - \overline{lpd_{-i}} \right) \tag{S34}$$
